# Biosynthesis of novel desferrioxamine derivatives requires unprecedented crosstalk between separate NRPS-independent siderophore pathways

**DOI:** 10.1101/2023.07.28.551008

**Authors:** Li Su, Yaouba Souaibou, Laurence Hôtel, Cédric Paris, Kira J. Weissman, Bertrand Aigle

## Abstract

Iron is essential to many biological processes, but its poor solubility in aerobic environments restricts its bioavailability. To overcome this limitation, bacteria have evolved a variety of strategies, including the production and secretion of iron-chelating siderophores. Here, we describe the discovery of four series of siderophores from *Streptomyces ambofaciens* ATCC23877, three of which are unprecedented. MS/MS-based molecular networking revealed that one of these series corresponds to acylated desferrioxamines (acyl-DFOs) recently identified from *S. coelicolor*. The remaining sets include unprecedented tetra- and penta-hydroxamate acyl-DFO derivatives, all of which incorporate a previously undescribed building block. Stable isotope labeling and gene deletion experiments provide evidence that biosynthesis of the acyl-DFO congeners requires unprecedented crosstalk between two separate NRPS-independent siderophore (NIS) pathways in the producing organism. The new derivatives, whose biological role(s) remain to be elucidated, not only illustrate the unanticipated biosynthetic potential of *S. ambofaciens*, but have interest in immuno-PET imaging applications.

**Importance:** Iron-chelating siderophores play important roles for their bacterial producers in the environment, but they have also found application in human medicine both in iron chelation therapy to prevent iron overload, as well as in advanced imaging applications. In this study we report the discovery of three novel series of related siderophores, whose biosynthesis depends on the interplay between two NRPS-independent (NIS) pathways in the producing organism *S. ambofaciens* – the first example to our knowledge of such functional cross-talk. We further reveal that two of these series correspond to acyl-desferrioxamines which incorporate four or five hydroxamate units. Although the biological importance of these novel derivatives is unknown, the increased chelating capacity of these metabolites may find utility in diagnostic imaging (for instance ^89^Zr based immuno-PET imaging) and other applications of metal chelators.

## Introduction

Although iron is abundant in the biosphere, its bioavailability is low due to the poor solubility of ferric oxide and hydroxide complexes. To obtain this essential nutrient, many bacteria synthesize and excrete specialized metabolites called siderophores which are capable of scavenging iron from their surroundings. The resulting high-affinity Fe^3+^ complexes are then re-acquired by siderophore uptake systems, followed by release of iron into the cytoplasm in Fe^2+^ form where it can play essential roles in many biological processes (1). To date, substantial knowledge has been accumulated concerning the diversity of siderophore chemical structures, as well as the molecular mechanisms underlying their synthesis, export, uptake and regulation (2–5). Siderophores often function as virulence factors for bacterial pathogens (e.g. aerobactin in enteric bacteria (6)) by sequestering iron from the animal or plant host, but conversely, they can also play important roles in pathogen control (7). Siderophores have also been implicated in mediating interspecies interactions (8). For example, siderophores produced by one species can be expropriated by another species (‘siderophore piracy’), or can induce the production of endogenous siderophores under iron-limited conditions (9, 10). Together, these various modes of siderophore action help to shape competition for iron in the environment, and facilitate various types of social interactions in soil and marine bacterial communities (11).

Siderophores are selective for ferric iron, but can also bind a range of other metals including zinc, manganese, copper, nickel, gallium, and aluminum (12). They are classified into four basic types depending on the moiety involved in iron chelation: catecholate, hydroxamate, carboxylate and mixed-type siderophores (13) (Fig. S1). These diverse structures are assembled via two principal means: non-ribosomal peptide synthetases (NRPSs) (14) and NRPS-independent siderophore (NIS) pathways (15). Notably, the siderophore products of NIS pathways often contain diamine, citric acid, and dicarboxylic acid building blocks (Fig. S1).

The biosynthetic gene clusters (BGCs) encoding siderophores are widely distributed in actinomycetes and *Streptomyce*s in particular, allowing the bacteria to respond to resource competition within the complex microbial community. In this context, *Streptomyces coelicolor* A3(2) has been well-studied, and shown to produce coelichelin and desferrioxamines E (DFO-E) and B (DFO-B) under iron-deficient conditions (16). Desferrioxamine B is of clinical importance, as it is approved by the US Food and Drug Administration (FDA) for removal of excess iron from the body (chelation therapy). Coelichelin, a tetrapeptide siderophore, is encoded by a NRPS gene cluster (*cch*), while the NIS pathway responsible for DFOs-B and E comprises a locus of six genes (*desABCDEF*) (17) (Fig. 1A), with both clusters located on the chromosome.

**FIG 1.**
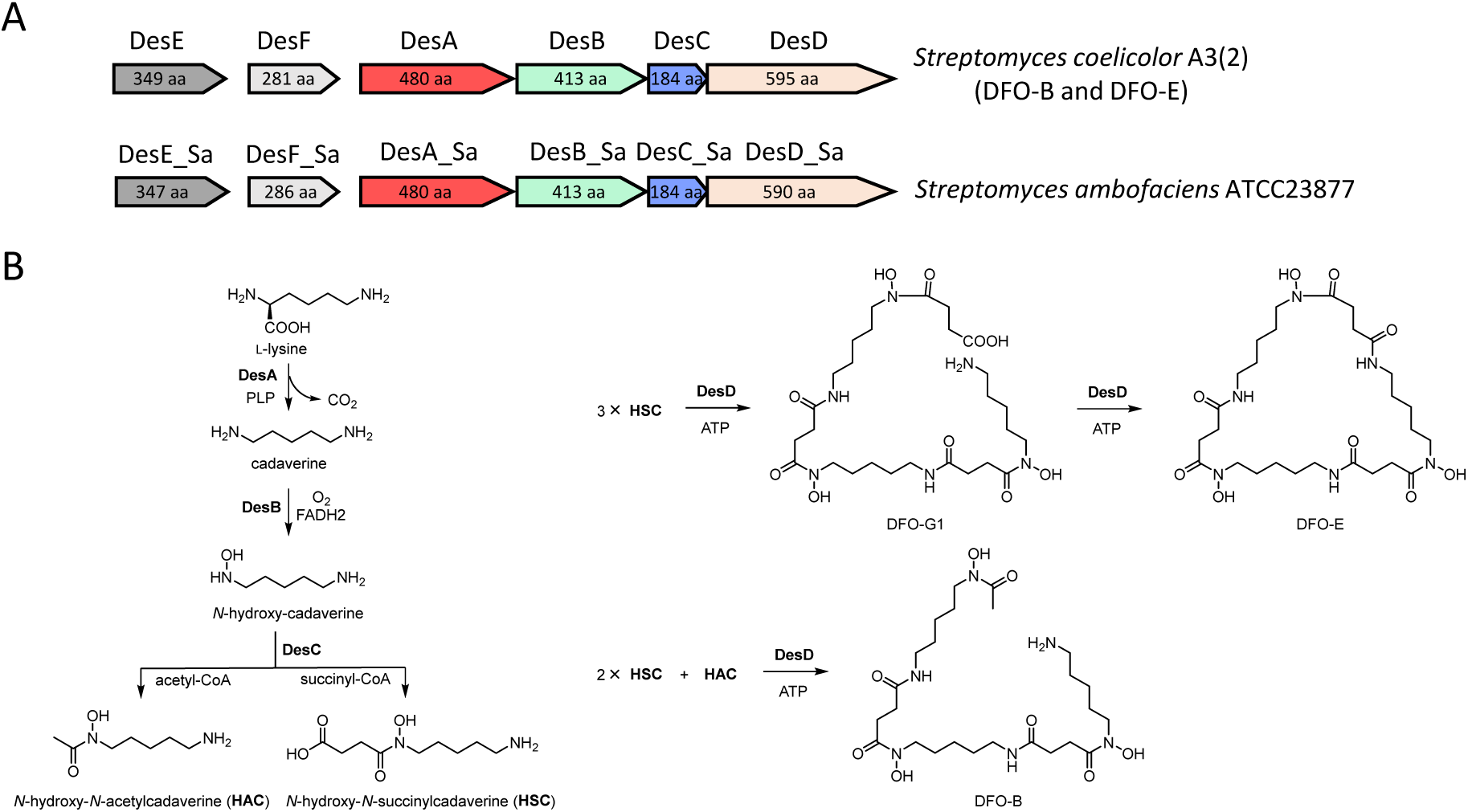
Representative biosynthetic gene clusters and pathway to NIS desferrioxamine. (A) The BGC for desferrioxamines (DFO-B and -E) in *S. coelicolor,* and the corresponding DFO BGC in *S. ambofaciens*. The same color indicates a shared enzymatic function, and the size of each protein in amino acids is indicated. (B) Currently proposed biosynthetic pathways to DFO-B and - E.

DFO-B and E are biosynthesized from L-lysine, acetyl-CoA and succinyl-CoA as building blocks (Fig. 1B). The first step involves generation of 1,5-diaminopentane (otherwise known as cadaverine) by decarboxylation of L-lysine, which is catalyzed by DesA, a pyridoxal 5-phosphate-dependent decarboxylase (16). This step is followed by mono-*N*-hydroxylation carried out by DesB, a FAD-dependent amine monooxygenase, resulting in *N*-hydroxy-cadaverine. This compound is then *N*-acetylated by DesC, an acyl CoA-dependent acyl transferase, to yield *N*-hydroxy-*N*-acetylcadaverine (HAC). Alternatively, DesC catalyzes *N*-succinylation to give *N*-hydroxy-*N*-succinylcadaverine (HSC). Three units of HSC are activated and condensed by DesD, an NRPS-independent peptide synthetase, to produce desferrioxamine G1 (DFO-G1) (18), followed by a ring-closing reaction which yields DFO-E (Fig. 1B). Linear DFO-B is composed of two units of HSC and a unit of HAC (Fig. 1B). The two other genes in the *des* cluster (Table S1), d*esE* and *desF*, encode respectively a cell surface-associated lipoprotein receptor component of an ABC transporter involved in uptake of the iron-chelated forms (called ferrioxamines), and a ferrioxamine reductase, an enzyme that removes iron from hydroxamate siderophore complexes.

Previous studies revealed that *S. ambofaciens* ATCC23877 harbors a *des* BGC and a *cch* BGC, both of which show strong conservation with the corresponding clusters in *S. coelicolor* (19, 20). The six enzymes encoded by the *S. ambofaciens desABCDEF* exhibit 83−95% amino acid sequence identity to their homologs in *S. coelicolor* (Fig. 1A, Table S1), and are responsible for the biosynthesis of DFO-B and E (20). In this work, we report four series of siderophores, in addition to the previously identified DFO-B and E, produced by both *Streptomyces ambofaciens* and mutants obtained by genetic engineering of the stambomycin gene cluster (21). One of these series corresponds to the acyl-DFOs, a class of recently-identified siderophores produced by *S. coelicolor* upon interaction with other actinomycetes (22). The additional three sets of DFOs, which we term acyl-DFOs+30, acyl-DFOs+200+30 and acyl-DFOs+200+200+30, were observed exclusively in *S. ambofaciens*. The numbers (+30, +200+30 and +200+200+30) in the named sets of acyl-DFOs represent the difference in mass units observed in MS/MS analysis. Results obtained from stable isotope labeling and gene deletion experiments allow us to propose a biosynthetic pathway for these novel acyl-DFO derivatives, which involves unprecedented crosstalk between separate NIS pathways. We further provide evidence that the ‘acyl-DFOs+200+30’ and ‘acyl-DFOs +200+200+30’ variants represent respectively tetra- and penta-hydroxamate structures distinct from those of the common bis-hydroxamate (e.g. bisucaberin (23)) and tris-hydroxamate iron (e.g. DFOs) chelators, and thus suggesting that NIS pathways are capable of extending oligomerization. Given the expanded binding capacity offered by the tetra- and penta-hydroxamate structures, the discovered metabolites may find utility in diverse applications as metal chelators (24).

## Results

### Identification of novel forms of desferrioxamines from *Streptomyces ambofaciens*

In the course of HPLC-MS analysis of a *S. ambofaciens* ATCC23877 mutant (K7N1/OE484) obtained by genetic engineering of the stambomycin gene cluster (21), four series of related, but previously undetected metabolites were identified as monoprotonated ions ([M+H]^+^) with *m*/*z* = 673, 687, 701, 715, 729, 743, 757 (first series); 703, 717, 731, 745, 759, 773, 787 (second series); 903, 917, 931, 945, 959, 973, 987 (third series); and 1103, 1117, 1131, 1145, 1159, 1173, 1187 (fourth series) (Fig. S2 and Table S2). The first two series of masses are separated from each other by 30 mass units and thus they were initially named ‘NMs’ (<u>N</u>ewly-detected <u>M</u>etabolite<u>s</u>) and ‘NMs+30’, respectively. The last two clusters of masses differ sequentially from NMs+30 by respectively +200 and +200+200, and as such were annotated as ‘NMs+200+30’ and ‘NMs+200+200+30’. Based on the observed masses, we originally hypothesized that the NMs could correspond to shunt polyketide products released from module 13 of the stambomycin polyketide synthase (PKS) (21), followed by the loss of a water molecule, either by dehydration or cyclization (the predicted cyclic form is shown in Fig. S2C) (25). However, these compounds were also identified in the wild type strain, albeit at lower levels, an observation which is inconsistent with classical PKS function (26) (Fig. S2B). Taken together, these data implied that the observed series of masses were unrelated to the stambomycin pathway.

Each mass series comprises 7 members, wherein each differs sequentially by +14 from its predecessor. In addition, the retention time for each NM/NM+30/NM+200+30/NM+200+200+30 group is quite close (Fig. S2). Both of these observations are consistent with compounds of related structure. However, on the basis of the integrated EIC peaks, we observed that the yields of the various series of metabolites differed both within and between the various strains, with the titers of NMs and NMs+30 superior to those of NMs+200+30 and NMs+200+200+30, and the high-molecular weight series present at 22−170% relative yield compared to the NMs (Fig. S2 and Table S3).

Using the Global Natural Products Social Molecular Networking platform (GNPS) (27), a molecular network of mutant K7N1/OE484 based on the fragmentation spectra was generated (Fig, S3). Via this method, compounds represented by nodes in the network are sorted based on the similarity of their fragmentation spectra. Nodes representing highly similar spectra are connected by edges and are likely to be structurally similar. Notably, the molecular network revealed that the NM series of compounds (shown with blue dots in Fig. 2 and Fig. S3) corresponds to various acyl-desferrioxamine siderophores (acyl-DFOs). Based on the GNPS ID hits, we identified the blue dots as C9−C15 acyl-DFOs (Fig. 2); although longer acyl-DFOs (C16 and C17) have been reported (22), they were not detected in this work. In addition, the corresponding ferrioxamines (acyl-FOs, Fig. 2 and Fig. S3), were also observed and linked to DFOs in the same subnetwork. Bisucaberin (23), DFO-E, DFO-D2 (28, 29), Desf-07 (29), and aluminum (Al)-chelated DFO-B were also annotated within this subnetwork, as were certain members of NMs+30, NMs+200+30 and NMs+200+200+30 (Fig. 2 and Fig. S3), consistent with their membership in the same metabolic family. The NM nodes are connected to DFOs and acyl-DFOs but do not correspond to compounds indexed within the GNPS database, identifying them as potentially novel acyl-DFO derivatives. We therefore re-designated them as acyl-DFOs+30, acyl-DFOs+200+30 and acyl-DFOs+200+200+30, respectively (Fig. 2 and Fig. S3).

**FIG 2.**
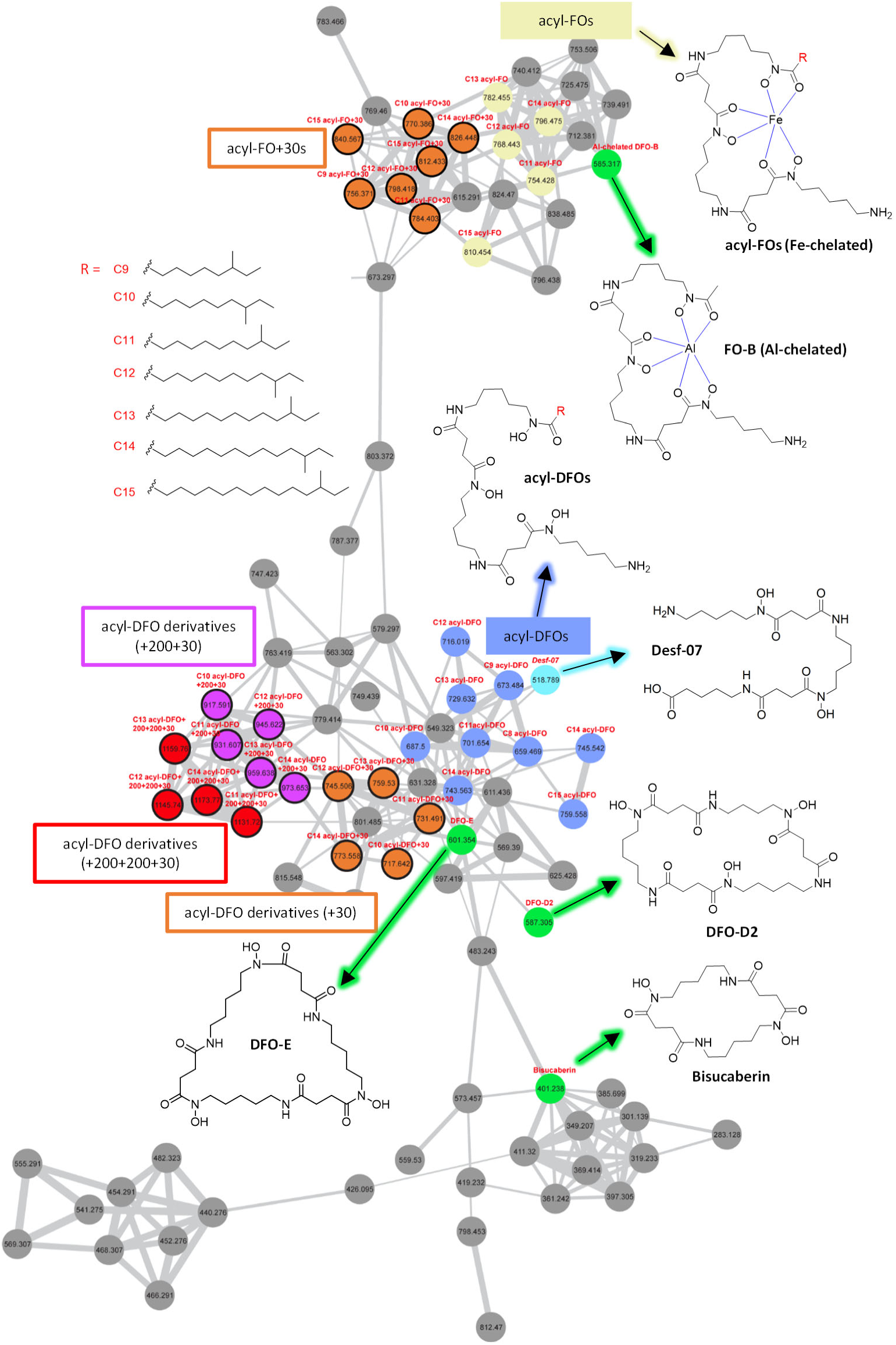
Annotation of the subnetwork related to the DFOs. The *m*/*z* value of the protonated parental ion ([M+H]^+^) is specified for each dot. The names of all known compounds are indicated above each dot, and their structures shown. The remaining members of acyl-DFOs+30 (C9 and C15), acyl-DFOs+200+30 (C9 and C15) and acyl-DFOs+200+ 200+30 (C9, C10 and C15) have not been presented in the network due to their relatively low parental ion densities (as shown in Fig. S2) and the subsequent lack of available MS/MS fragments in MS/MS full scan model. The complete molecular network is shown in Fig. S3.

### Structure analysis of novel acyl-DFO derivatives

As the relatively low yields of the novel metabolites compounded with the difficultly of chromatographic separation of closely-similar structures (Fig. S2) precluded structure elucidation by NMR, we opted for an MS/MS-based analysis. As a starting point, inspection of the previously identified MS/MS fragments for DFOs and acylated DFOs (22), revealed that certain fragments were common to all 7 acyl-DFOs (C9–C15) (e.g. *m*/*z* [M+H]^+^ = 401, 319, 201) (Fig. S4). Thus, these fragments must arise from the shared amine portion of the structures. We also observed fragments encompassing the acyl regions, as these differed by +14 between the acyl-DFO family members consistent with the presence of additional -CH_2_ units (e.g. the series *m*/*z* [M+H]^+^ = 397, 515 and 597 arising from C12 acyl-DFO vs. 411, 529, and 611 from C13 acyl-DFO) (Fig. S4). In the case of the novel derivatives, we observed the same acyl-derived fragments as for the corresponding acyl-DFOs, allowing in each case identification of the acyl group appended to the DFO+30, DFO+200+30 and DFO+200+200+30 series of compounds (Fig. 3 and Fig. S4–7). In contrast, a new set of fragments common to all of the novel metabolites was identified with masses of +30 relative to those observed in DFOs and acyl-DFOs (*m*/*z* [M+H]^+^ = 431, 349, 231; Fig. 3 and Fig. S4–7). Thus, the +30 region is located within the opposite region of the compounds to the acyl terminus.

**FIG 3.**
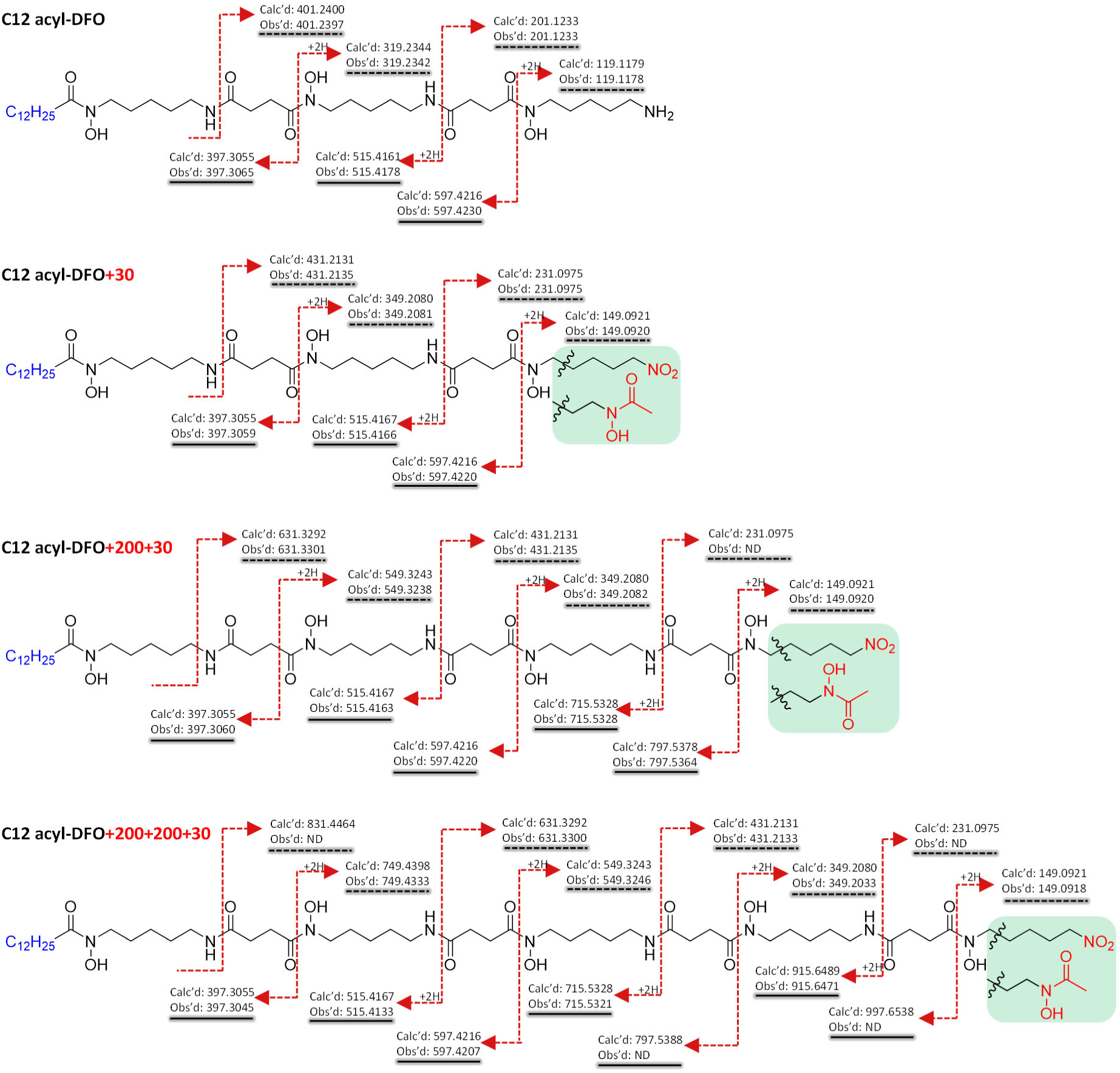
Comparison of the observed HPLC-MS/MS fragmentation patterns of C12 acyl-DFO and its congeners. The structure and MS/MS fragment assignment of C12 acyl-DFOs are based on previously-published work (22). Comparison with the MS/MS data obtained on C12 acyl-DFO+30, C12 acyl-DFO+200+30 and C12 acyl-DFO+200+200+30 reveals a series of shared fragments (solid underline), as well as divergent fragments (dashed underlines) characteristic of the extended structures. The structure of the portion of the molecules highlighted in green has not been conclusively determined, but the available data are most consistent with the *N*-hydroxy, *N*-acetyl version.

Detailed analysis of the fragmentation patterns of the acyl-DFOs+200+30 and acyl-DFOs+200+200+30 also allowed us to propose that the +200 and +200+200 arise via incorporation of one or two additional building blocks of HSC, respectively, into the metabolite backbones (Fig. 4). For example, fragments encompassing the acyl group of C12 acyl DFO+30 (*m*/*z* [M+H]^+^ = 515 and 597) are additionally observed at +200 in C12 acyl-DFO+200+30 (*m*/*z* [M+H]^+^ = 715 and 797) (Fig. 3), while these same +200 fragments as well as +200+200 fragments (*m*/*z* [M+H]^+^ = 915 and 997) were seen from C12 acyl-DFO+200+200+30 (Fig. 3 and Fig. S6, 7). Additional sets of fragments show this pattern: *m*/*z* [M+H]^+^ = 349 and 431 common to all acyl-DFOs+30 can be correlated with +200 fragments in both the acyl-DFO+200+30 and acyl-DFO+200+200+30 series (*m*/*z* [M+H]^+^ = 549 and 631), while the corresponding +200+200 fragments are observed with the acyl-DFOs+200+200+30 (*m*/*z* [M+H]^+^ = 749 and 831) (Fig. 3 and Fig. S6, 7). The +200 and +200+200 versions of DFO-B, DFO-D2, DFO-E and DFO-G1 were not seen.

**FIG 4.**
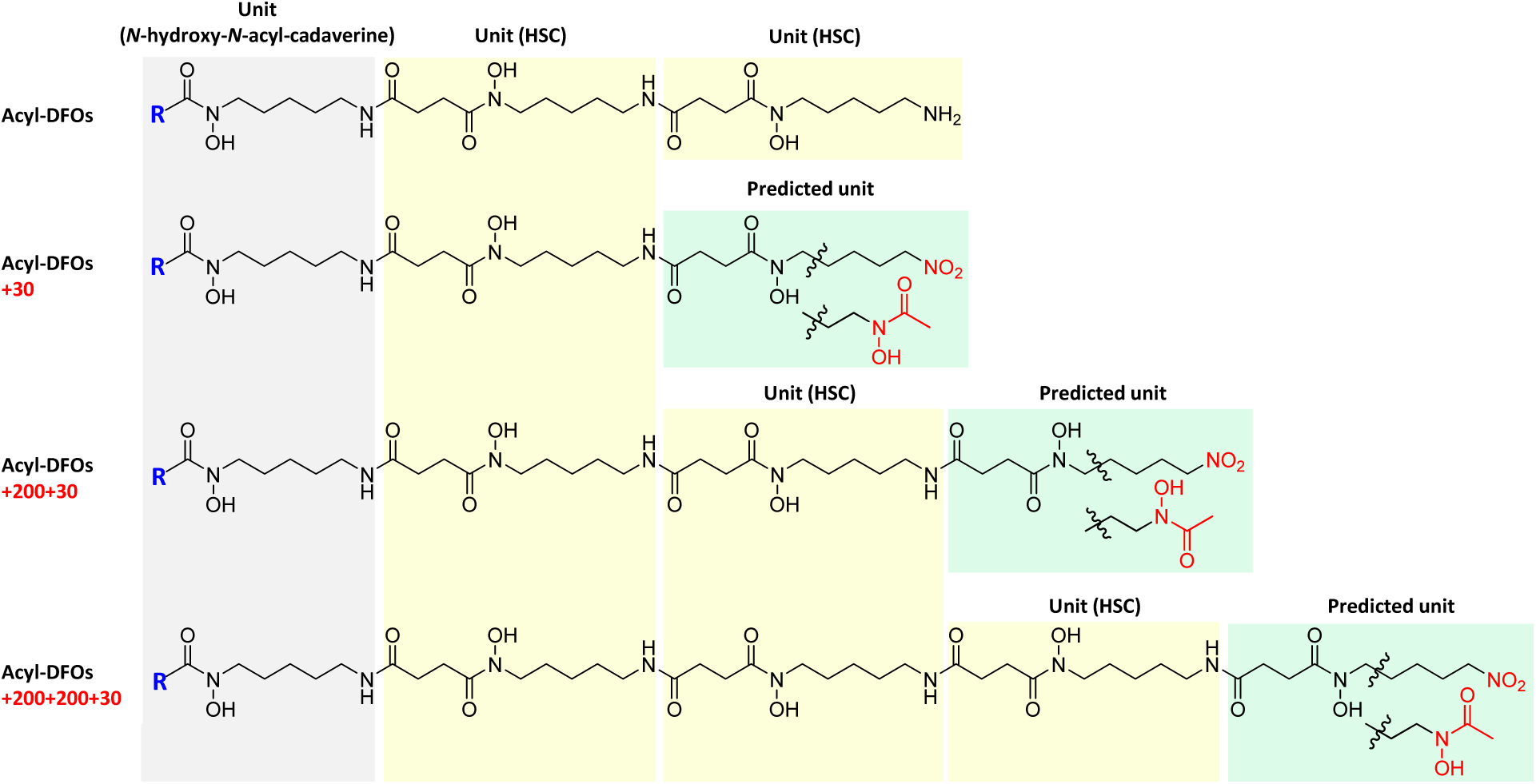
Comparison of the monomer composition of acyl-DFOs+30, acyl-DFOs+200+30 and acyl-DFOs+200+200+30. The structures of the acyl-DFOs were published previously (22), while the structures of the additional derivatives were predicted on the basis of the MS/MS fragmentation patterns. The building blocks shown in different colors are linked together by amide bonds to form the various DFOs (18, 52). The “+200” refers to incorporation of an additional HSC unit, while the “+30” is predicted to originate from one of two alternative modifications to the terminal amine group, with the *N*-hydroxy, *N*-acetyl congener being more likely.

Nonetheless, at this stage, the precise structural differences giving rise to the +30 mass discrepancy were unclear. One possible explanation we considered was transformation of the terminal amine group (−NH_2_) into a nitro group (−NO_2_). This chemistry has been reported to occur during LC-ESI-MS analysis, as electrochemical reactions can take place at the liquid-metal interface on the capillary, leading to reactive oxygen species (e.g. •OOH and H_2_O_2_) capable of amine oxidation (30, 31). Alternatively, the NO_2_ could derive from spontaneous enzymatic oxidation of NH_2_. To assess this latter possibility, we carried out a BLASTP search with *S. ambofaciens* using the sequence of PrnD from *Pseudomonas fluorescens*, which encodes a [2Fe-2S] Rieske non-heme *N*-oxygenase involved in the biosynthesis of the antifungal antibiotic pyrrolnitrin (32, 33). This analysis identified SAM23877_0811, with 39% amino acid sequence identity to PrnD (Fig. S8). The second overall explanation is that the +30 compounds incorporate an alternative 1,3-diaminopropane-derived building block (*vida infra*).

### Incorporation *in vivo* of isotopically-labeled lysine

In order to differentiate between the potential origins of the +30 moiety (an enzymatic or chemical derivative of lysine vs. integration of an alternative monomer) as well as to obtain more direct evidence for the number of HSC building blocks present in each series of molecules, we supplemented cultures of strain K7N1/OE484 with isotopically-labeled lysine (^13^C_6_, ^15^N_2_-lysine, 1 mM). If lysine were the origin of the +30 moiety, we would have expected to observe as many as three Lys molecules incorporated into the acyl-DFOs+30, four into the acyl-DFOs+30+200, and five into the acyl-DFOs+200+200+30. Instead, using the C12 acyl-DFO series as an example, analysis of the isotopic distribution of the parent ions in fed vs. unfed samples, showed that while the C12 acyl-DFO incorporated up to three units of labeled lysine (mass shifts of +7, +14 and +21 Da) (Fig. 5B), only a maximum of two lysines were integrated into the +30 derivative, three into the +200+30 compound and four into the +200+200+30 analogue (Fig. 5D, F, H). These data are fully consistent with the +200 masses arising from incorporation of additional HSC units, but not with *N*-oxidized lysine as the origin of the +30 mass difference. Furthermore, integration of labeled lysine did not increase the mass of a characteristic fragment of C12 acyl-DFO+30 containing the +30 region (*m*/*z* [M+H]^+^ = 149), while a deletion mutant in the putative *N*-oxygenase SAM23877_0811 still produced the +30/+200+30/+200+200+30 congeners of the acyl-DFOs (Table S3).

**FIG 5.**
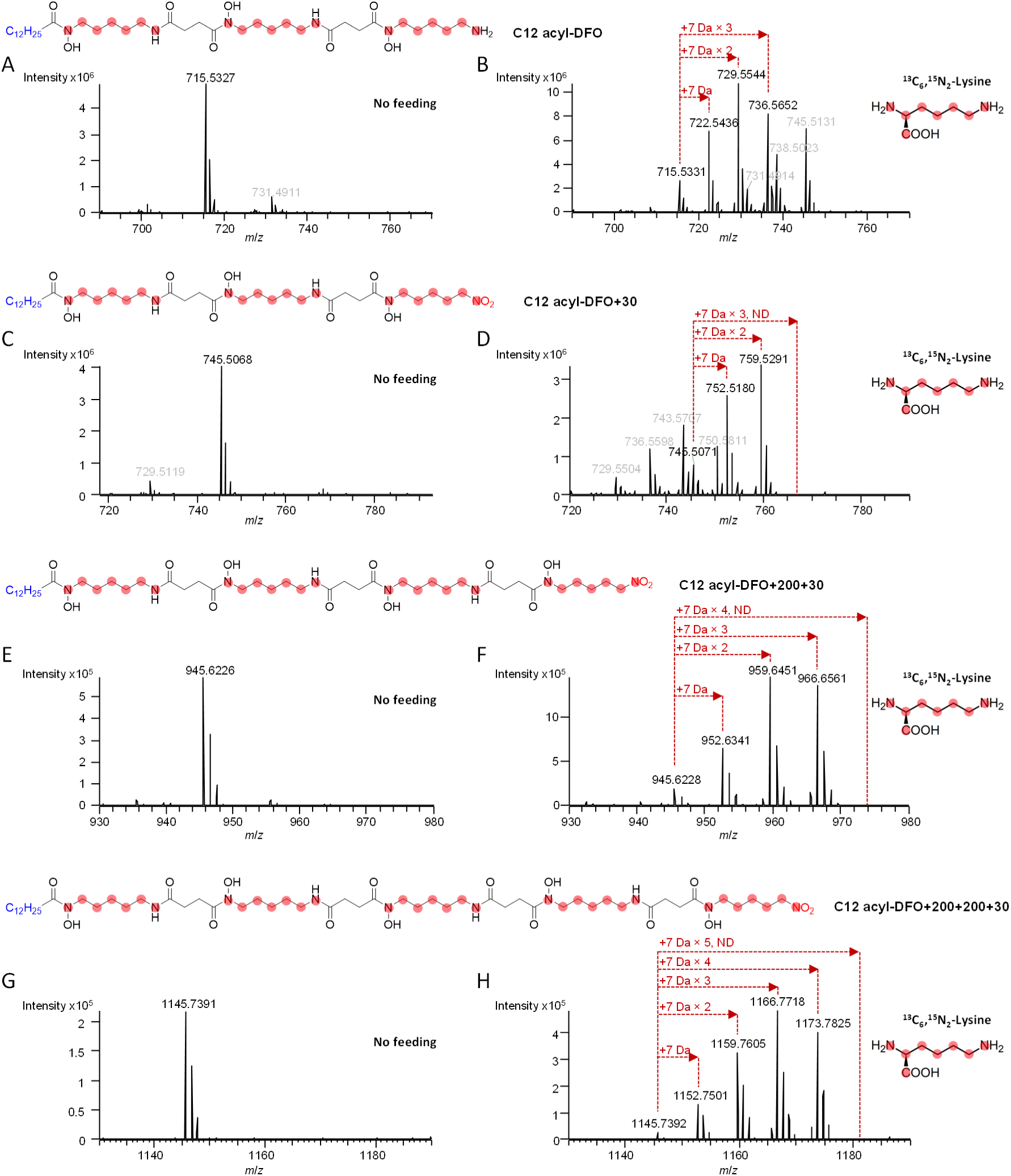
Patterns of isotope incorporation into C12 acyl-DFO and its three novel derivatives in the absence (A, C, E, G) and presence (B, D, F, H) of fed ^13^C6, ^15^N2-lysine. The mass spectrum of the parent ion (*m*/*z* [M+H]^+^) is shown in each case. The expected labeling patterns are shown as red dots on the structures. The absence of +7 Da × 3, +7 Da × 4 and +7 Da × 5 peaks in D, F and H, respectively, excluded the possibility the compounds incorporate a Lys-derived nitrosylated building block (Fig. S8). C11 acyl-DFO+30 was also detected (*m*/*z* values shown in grey in panels A and B) in the spectrum of C12 acyl-DFO+30, as the two metabolites co-eluted (Fig. S2). Similarly, peaks indicated in grey in C and D correspond to C13 acyl-DFO. ND = not detected.

### A putative 1,3-diaminopropane-derived building block involved in the biosynthesis of novel DFO derivatives

In general, the structures of NIS-mediated siderophores, and in particular the hydroxamate-type, incorporate a diamine moiety, such as 1,3-diaminopropane, 1,4-diaminobutane (putrescine) or 1,5-diaminopentane (cadaverine) (Fig. S11). The origins of the backbones of 1,3-diaminopropane, putrescine and cadaverine are (*S*)-2-amino-4-oxobutanoic acid, L-ornithine and L-lysine, respectively. More specifically, in the biosynthetic pathway of schizokenin in *Nostoc sp. PCC 7120 (also known as Anabaena sp. PCC 7120)*, (*S*)-2-amino-4-oxobutanoic acid is transformed into 1,3-diaminopropane by sequential action of an L-2,4-diaminobutyrate:2-ketoglutarate 4-aminotransferase (DABA AT, *all0396*) and a L-2,4-diaminobutyrate decarboxylase (DABA DC, *all0395*) (Fig. 6A, B) (34–36). Genes encoding the DABA AT and DABA DC pair are also present in the BGCs of the 1,3-diaminopropane-based siderophores synechobactins (37), rhizobactin 1021 (38), and acinetoferrin (39).

**FIG 6.**
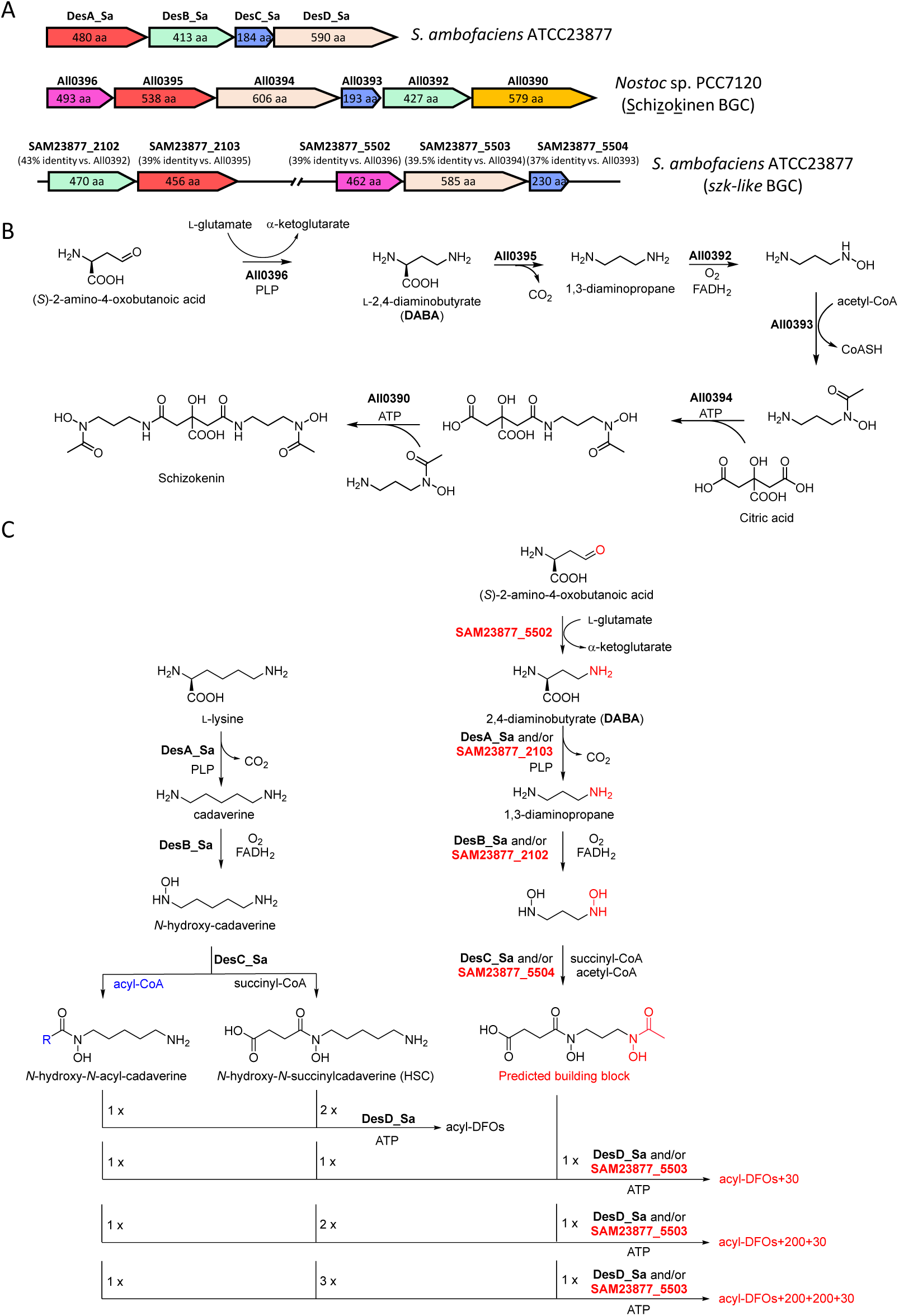
Predicted biosynthetic pathway to all DFO variants described in this study. (A) The genomic organization of the potential gene cluster involved in the biosynthesis of the predicted building block, relative to the enzymes involved in the biosynthesis of schizokenin in *Nostoc* sp. PCC7120 and DFOs in *S. ambofaciens*. Identities on the AA sequence level are shown. Enzymes All0391 (MFS transporter) and DesE and DesF are not shown. (B) Proposed biosynthetic pathway to schizokenin. (C) Proposed biosynthetic pathway to the acyl-DFOs+30, acyl-DFOs+200+30 and acyl-DFOs+200+200+30.

Interestingly, we identified two loci on the chromosome of *S. ambofaciens*, one of which contains homologs of *all0392* (43% identity at the AA sequence level) and *all0395* (39%) involved in the biosynthesis of schizokenin, while the other comprises homologs of *all0396* (39%), *all0393* (37%), and *all0394* (40%) (Fig. 6A). It thus appears that a portion of an ancestral *szk* cluster was relocated to an alternative region within the *S. ambofaciens* genome. Nonetheless, a homolog of gene *all0390* encoding the second NIS enzyme required for schizokenin biosynthesis is not present in either locus, and thus the pathway appears incomplete. In accord with this observation, no schizokenin was detected in cultures of *S. ambofaciens* wild type and its mutants (data not shown). Nonetheless, the availability of additional genes capable of diamine synthesis prompted us to consider a new pathway to a 1,3-diaminopropane-derived building block in *S. ambofaciens* (Fig. 6C), based on collaboration between enzymes encoded by the desferrioxamine and schizokenin-like (*szk*-like) gene clusters.

In this scheme, DesABC would be responsible for generation of the cadaverine-based units HSC and *N*-hydroxy-*N*-acyl-cadaverine, while either of the enzyme pairs SAM23877_5502 (DABA AT)/SAM23877_2103 (DABA DC) and/or SAM23877_5502 (DABA AT)/DesA_Sa (DABA DC) (35) would produce 1,3-diaminopropane. Subsequently, 1,3-diaminopropane would be *N*-hydroxylated by DesB and SAM23877_2102 (or *N*-hydroxylated twice by one of the two enzymes) on its two amino termini, followed by *N*-acetylation and *N*-succinylation by DesC and SAM23877_5504, respectively (or via bifunctional catalysis by one of the two enzymes). The final step would involve concatenation of the monomers via amide bonds, catalyzed by the two NIS synthetases DesD and SAM23877_5503 (or alternatively by one of the two), resulting in the synthesis of acyl-DFOs+30, acyl-DFOs+200+30 and acyl-DFOs+200+200+30.

Given that the +30 building block could in principle be incorporated into other desferrioxamines produced by the strain, we also scrutinized the extracts for the corresponding versions of DFO-B, DFO-E and DFO-G1. This analysis revealed the +30 forms of both linear DFO-B and DFO-G1 (Fig. S5), but not of cyclic DFO-D2 and DFO-E.

### Testing the biosynthetic hypothesis by gene deletion within the *des* and *szk*-like clusters

To evaluate whether the biosynthesis of novel DFO derivatives requires crosstalk between the *des* and *szk*-like clusters, several mutants were created using the stambomycin engineering mutant K7N1 as the base strain. We first deleted genes *desC* (*N*-acetylase/*N*-succinylase) and *desD* (NIS synthetase) in the *des* cluster using CRISPR-Cas9 (Fig. S12), followed by the introduction of plasmid pOE484 (in order to make it equivalent to the control strain K7N1/OE484), resulting in the mutant K7N1/Δ*desCetD*/OE484. Analysis by HPLC-MS of the mutant relative to the parental strain revealed that all DFO production was completely abolished (Fig. S15B). Thus, the *des* cluster is essential for biosynthesis of all types of DFOs in *S. ambofaciens*. The split *szk*-like cluster was also inactivated by CRIPSR-Cas9-mediated deletion by generation of a mutant in each locus, K7N1/Δ*sam2102et03*/OE484 (*N*-hydroxylase and DABA DC mutant) and K7N1/Δ*sam5502et03et04*/OE484 (DABA AT, NIS synthetase and *N*-acetylase/*N*-succinylase mutant) (Fig. S13, 14). Although neither *szk* deletion mutant completely abrogated DFO production (Fig. S15C, D), the yields of the various series of DFOs were all reduced by 10–100 fold relative to K7N1/OE484 (Table S3), strongly implicating the *szk*-like cluster in their biosynthesis (Fig. 6C).

Indeed, the continued production of these novel acyl-DFOs by K7N1/Δ*sam2102et03*/OE484 is fully consistent with the proposed functional redundancy between the *N*-hydroxylases SAM23877_2102 and DesB, and the DABA DCs SAM23877_2103 and DesA_Sa. Production of +30 containing acyl-DFOs by mutant K7N1/Δ*sam5502et03et04*/OE484 was less readily explained, as no enzyme encoded by the *des* cluster could functionally complement the missing SAM23877_5502 for DABA synthesis. However, reinspection of the *S. ambofaciens* genome sequence revealed multiple SAM23877_5502 homologs which could rescue the missing activity, including SAM23877_1053 (32% identity and 41% similarity at the aa level), SAM23877_6419 (29% identity, 42% similarity), and SAM23877_1939 (28% identity, 41% similarity). SAM23877_1939 is especially interesting in this context, as it functions as a DABA AT (EctB) during biosynthesis of DABA in the pathway to ectoine and hydroxyectoine (Fig. S16) (40).

### *In vivo* incorporation of labeled acetate

In our proposed pathway to the alternative building block (Fig. 6), a 1,3-diaminopropane-derived unit bearing both an *N*-succinyl and an *N*-acetyl is generated via *szk*-like chemistry. As acetate is the distinguishing feature of this monomer relative to *N*-hydroxy-*N*-acyl-cadaverine and HSC, we reasoned that feeding of 2-^13^C-acetate would result in more pronounced labeling of all the +30 compounds relative to the acyl-DFOs, while MS/MS analysis would help us to localize the sites of increased incorporation. It is important to note that we did not expect exclusive labeling of the +30 metabolites, as 2-^13^C-acetate could be converted to succinate by the TCA cycle as well as into long-chain fatty acids, and thus incorporated into the acyl-DFOs.

Compared to the K7N1/OE484 strain cultivated in unsupplemented medium, the +1 peak for C12 acyl-DFO following growth in the presence of 1 mM sodium acetate-2-^13^C, increased in relative intensity from 41.6% (Fig. 7A) to 52.5% (Fig. 7B). Likewise, the +1 peaks for DFO-B and G1 increased in relative intensity from 28.4% to 34.9%, and from 31.3% to 38.2%, respectively (Table S7). These observations are in accord with the conversion of the fed acetate into more elaborate precursors followed by incorporation. However, the intensity of the +1 peak for the C12 acyl-DFO+30 increased by an additional 4.43% compared to that of C12 acyl-DFO (from 52.5% in Fig. 7B to 56.9% in Fig. 7D) while the +2-peak showed similarly higher enrichment, consistent with greater incorporation of acetate-2-^13^C into C12 acyl-DFO+30 relative to C12 acyl-DFO (Fig. 7 and Table S7).

**FIG 7.**
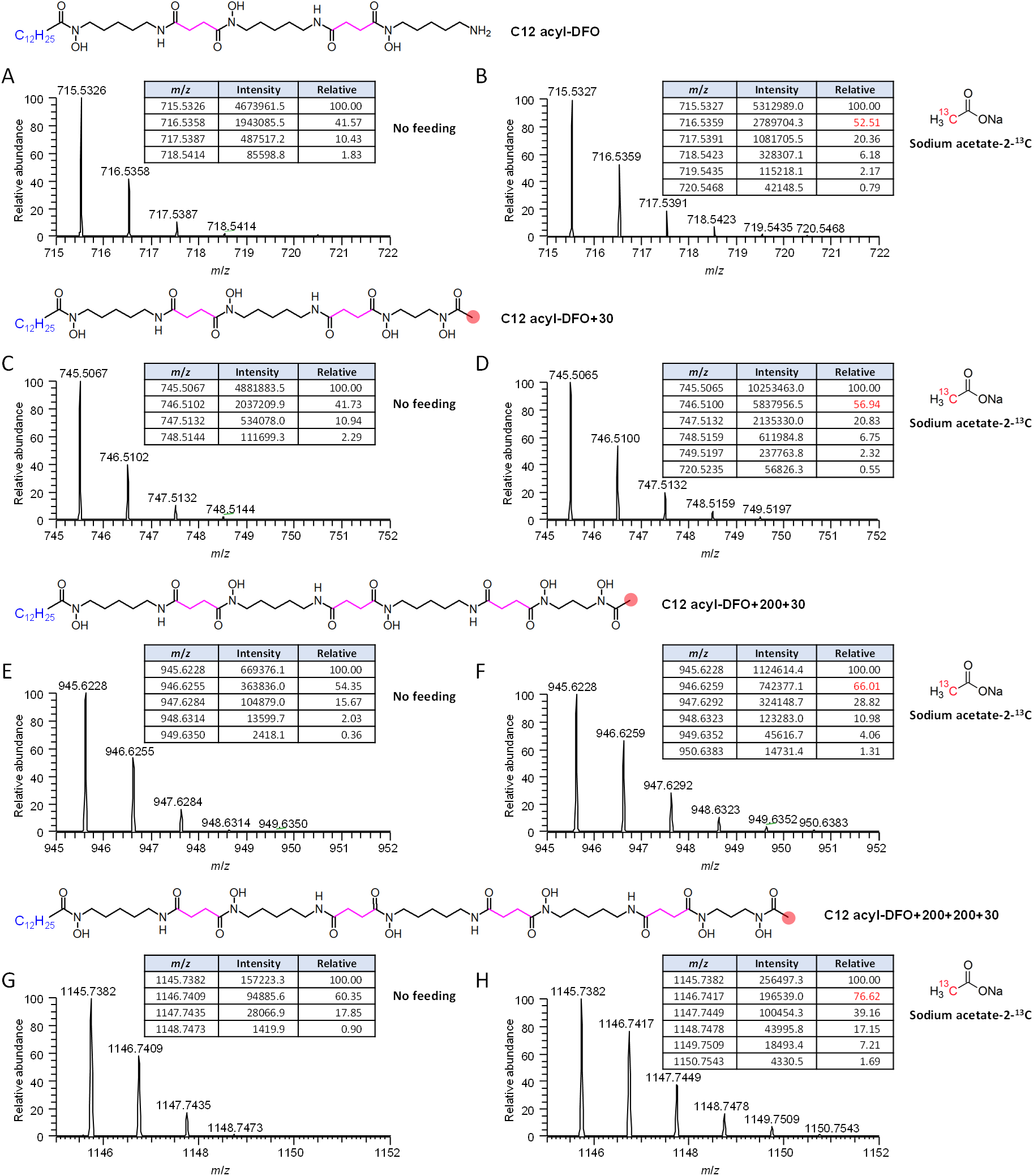
Patterns of isotope incorporation into C12 acyl-DFO and its three novel congeners in the absence (A, C, E, G) and presence (B, D, F, H) of fed sodium acetate-2-^13^C. The expected site of labeling of the proposed alternative building block is shown as a red dot, and the observed relative intensity of the +1 peaks highlighted in red. Succinate units, which can also incorporate fed acetate, are indicated in pink. The complete data set for all DFOs (C9–C15) are included in Table S7.

In order to track the site of incorporation, we sought to compare the MS/MS fragmentation patterns between C12 acyl-DFO and C12 acyl-DFO+30. For this, we relied on fragments of the molecules which are diagnostic for their structures, as they exhibit a mass difference of +30 and so encompass the divergent regions: *m*/*z* [M+H]^+^ = 319 and 119 for acyl-DFOs, versus *m*/*z* [M+H]^+^ = 349 and 149 for acyl-DFOs+30 (Fig. 3). This analysis showed that the +1 peaks of 319 and 119 were observed at 54.72% and 6.04% relative intensity, respectively, in the presence of 2-^13^C-acetate. As a +1 peak of 5.35% intensity would be expected for the 119-fragment based on natural ^13^C abundance (5C × 1.07%), these data show that intact acetate incorporation into this portion of C12 acyl-DFO was minimal (Fig. 8A). In contrast, the +1 peaks of the 349 and 149 fragments were of substantially higher intensity (60.63% and 10.52%, respectively) (Fig. 8A), providing evidence that they both incorporate an intact acetate unit. Thus overall, the acetate feeding experiments are in line with our proposed biosynthetic pathway (Fig. 6).

**FIG 8.**
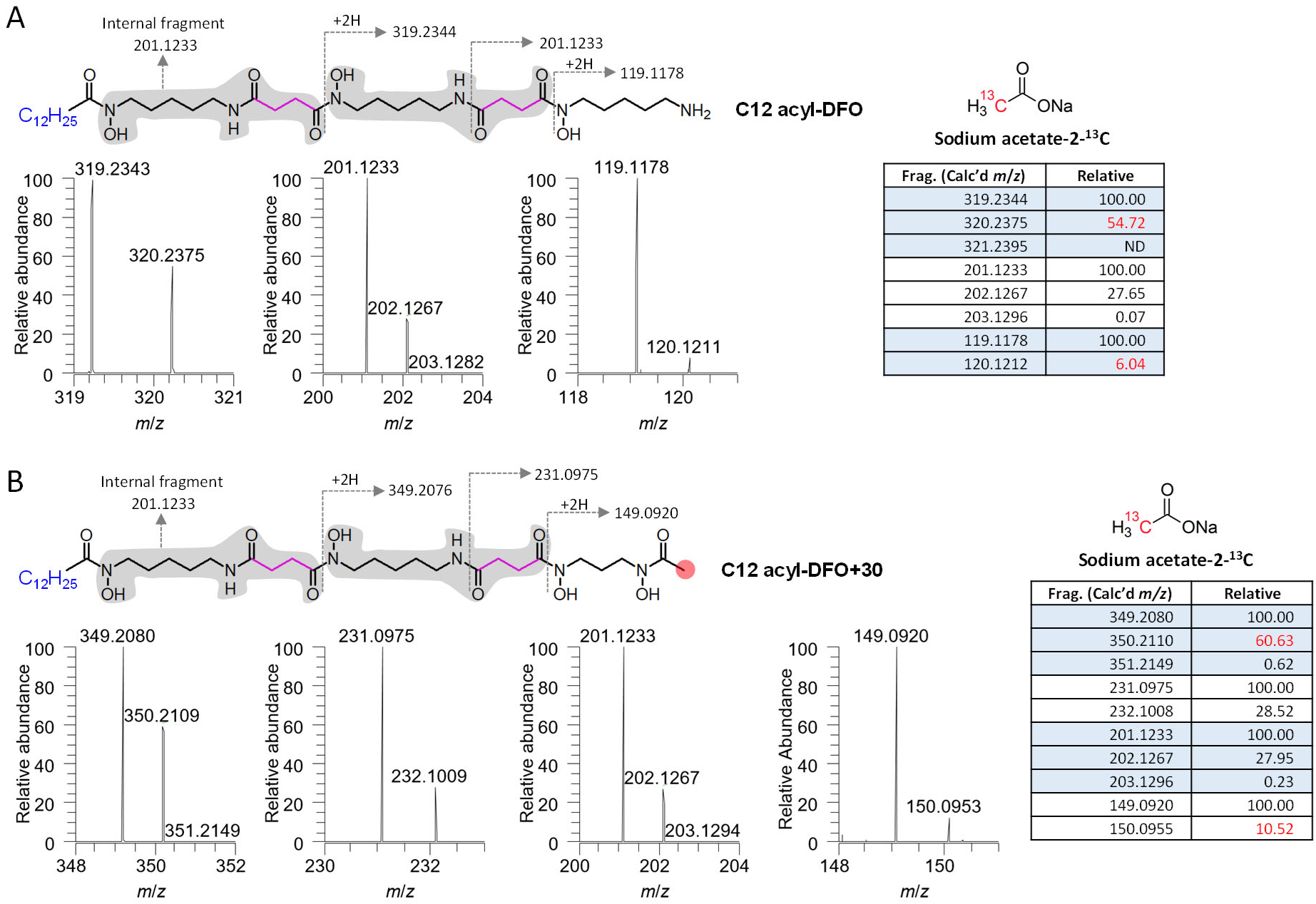
Representative MS^2^ fragments of C12 acyl-DFO and C12 acyl-DFO+30 after feeding with sodium acetate-2-^13^C. The indicated fragments derive from parental ions 716.6359 and 746.5100 in Figs. 7B and 7D, respectively. We chose to focus our analysis on the *m*/*z* [M+H]^+^ = 349/319 and 149/119 pairs, as the origin of the 201/231 fragments is ambiguous (terminal vs. internal fragment in C12 acyl-DFO). ND = not detected. The complete data sets are provided in Table S8.

## Discussion

Acyl-DFOs are desferrioxamines bearing a saturated fatty acid of variable length (C9−C17) on the terminal hydroxamate group (22). While such amphiphilic siderophores are commonly produced by marine bacteria (e.g. marinobactins (41), aquachelins (41), amphibactins (42), ochrobactins (43), and synechobactins (37)), they are rare in terrestrial bacteria such as *S. ambofaciens*. A potential explanation for this disparity is the ability of such metabolites to embed themselves in the cell membrane (acyl chain lengths from C14−C21; (44)), or to form micelles and vesicles (C12−C16 (42)) – features that should help limit diffusion into the open ocean from the bacterial cells. The physical structure of soil that intrinsically limits the diffusion of molecules was then thought to account for their relative scarcity in terrestrial bacteria (45). Our finding that in addition to *S. coelicolor*, the soil bacterium *S. ambofaciens* (41) also produces acyl-DFOs, rather argues for the commonality of DFO composition between soil and marine bacteria.

Nonetheless, we have identified three novel sets of related metabolites in *S. ambofaciens*, acyl-DFOs+30, acyl-DFOs+200+30, and acyl-DFOs+200+200+30, which are not present in *S. coelicolor*, and provided evidence for their synthesis via an unprecedented interplay between two distinct NRPS-independent siderophore pathways. An incomplete *szk*-like pathway is also present in *S. coelicolor* suggesting that it should also have the ability to produce these novel acyl-DFO derivatives at least under certain growth conditions. Indeed, acyl-DFO production from *S. ambofaciens* is environment-dependent, with yields substantially higher when the strain is grown on liquid M5 medium (21) instead of 26A agar medium ((9), data not shown), reflecting the fact that nutrients and precursors present under different cultivation conditions can influence the product spectrum within a particular metabolite family. Of note, we also observed the +30 forms for DFO-B and DFO-G1 (Fig. S5), which demonstrates that the interplay between the *des* and the *szk*-like pathways is not restricted to the acyl-DFOs, but encompasses all linear desferrioxamine derivatives. Future work will focus on elucidating the biological functions of these compounds, particularly with regard to their metal-chelating capacity.

In conclusion, we note that the ability of DFO-B to chelate [^89^Zr]Zr^4+^ has been exploited clinically for immuno-PET imaging (46). However, the complex exhibits suboptimal stability *in vivo*, as DFO-B only offers six coordination sites for Zr^4+^ via its three hydroxamate moieties (46). This observation has inspired the chemical synthesis of various octadentate versions of DFO-B capable of saturating the Zr^4+^ coordination sphere (47, 48), including analogs exhibiting improved water solubility (49). Our results show that nature also assembles a range of extended DFOs, which may find utility in diagnostic imaging and other applications of metal chelators (24). Given the evident difficulty of purifying the metabolites from the native host, however, it would be worthwhile to chemically synthesize the compounds (50) for both biophysical characterization and elucidation of their biological roles.

## Materials and Methods

### Materials

All reagents and chemicals were obtained from Sigma-Aldrich, except the following: BD (tryptone, yeast extract, TSB powder), Thermo Fisher Scientific (Tris), VWR (glycerol, NaCl, NaNO_3_), ADM France (NutriSoy flour), and New England Biolabs (T4 DNA ligase, restriction enzymes). Oligonucleotide primers were synthesized by Sigma-Aldrich. DNA sequencing of PCR products was performed by Sigma-Aldrich. PCR reactions were carried out with Taq DNA polymerase (Thermo Fisher Scientific), or Phusion High-Fidelity DNA polymerase (Thermo Fisher Scientific) when higher fidelity was required. Isolation of DNA fragments from agarose gel, purification of PCR products and extraction of plasmids were carried out using the NucleoSpin® Gel and PCR Clean-up or NucleoSpin® Plasmid DNA kits (Macherey Nagel, Hoerdt, France).

### Strains and media

Unless otherwise specified, all *Escherichia coli* strains were cultured in LB medium (yeast extract 10 g, tryptone 5 g, NaCl 10 g, distilled water up to 1 L, pH 7.0) or on LB agar plates (LB medium supplemented with 20 g/L agar) at 37 °C. *Streptomyces ambofaciens* ATCC23877 and the mutants thereof were grown in TSB (TSB powder 30 g (tryptone 17 g, soy 3 g, NaCl 5 g, K_2_HPO_4_ 2.5 g, glucose 2.5 g), distilled water up to 1 L, pH 7.3) or on TSA plates (TSB medium supplemented with 20 g/L agar), and sporulated on SFM agar plates (NutriSoy flour 20 g, D-mannitol 20 g, agar 20 g, tap water up to 1 L) at 30 °C. All strains were maintained in 20% (*v*/*v*) glycerol in 2 mL Eppendorf tubes and stored at −80 °C. For fermentation of *S. ambofaciens* ATCC23877 and its mutants, spores were streaked on TSA with appropriate antibiotics (apramycin, etc..) and after incubation for 48 h at 30 °C, a loop of mycelium was used to inoculate 7 ml of MP5 medium (yeast extract 7 g, NaCl 5 g, NaNO_3_ 1 g, glycerol 36 mL, MOPS 20.9 g, distilled water up to 1 L, pH 7.4) supplemented with selective antibiotics and sterile glass beads, followed by incubation at 200 rpm and 30 °C for 24–48 h. Finally, the seed culture was centrifuged and resuspended into 2 mL fresh MP5 before being inoculated into 50 mL MP5 medium in a 250 mL Erlenmeyer flask, and cultivated at 200 rpm and 30 °C for 4 days. Labeled molecules were fed to the media at a final concentration of 1 mM after inoculation of seed culture at 24h incubation.

### CRISPR-Cas9 mediated genetic engineering

Plasmid pCRISPomyces-2 was used for construction of all mutants generated in this study (Tables S4, 5). For each gene or locus to be deleted, the crRNA sequence was selected to match the DNA segment which contains NGG on its 3′ end (N is any nucleotide, and the NGG corresponds to the protospacer-adjacent motif (PAM)). The annealed crRNA fragment and two homologous arms (HAL and HAR, flanking the target region) were sequentially inserted into the delivery plasmid pCRISPomyces-2 using the restriction sites *Bbs*I and *Xba*I, respectively, to afford the specific recombinant plasmid for each mutant (Fig. S12–14).

### LC-ESI-HRMS analysis of fermentation metabolites

The fermentation broth was centrifuged at 4,000 x *g* for 10 min. The acyl-DFOs and their derivatives were then extracted from the mycelia by first resuspending the cells in 40 mL distilled water, followed by centrifugation (4,000 × *g*, 10 min, repeated 3×) to remove the water-soluble components. After decanting the water, the cell pellets were weighed and extracted with MeOH by shaking at 150 rpm for 2 h at room temperature. Thereafter, the MeOH extracts were filtered to remove the cell debris, followed by rotary evaporation to dryness. The obtained extracts were then dissolved in MeOH, whose volume was determined according to the initial weight of the mycelia (70 μL MeOH to 1 g of cell pellet). The resulting mycelium extracts were then centrifuged at 16,000 x *g* at 4 °C for 20 min and analyzed in heated positive electrospray mode (HESI+) by HPLC-HRMS on a Thermo Scientific Orbitrap ID-X Tribrid Mass Spectrometer using an Alltima™ C18 column (2.1 × 150 mm, 5 μm particle size, Grace-Alltech) or an Acclaim™ C18 column (2.1 × 100 mm, 2.2 μm particle size, ThermoScientific). Separation was carried out with ultrapure water (Purelab Flex system, Veolia) containing 0.1% formic acid (A) and acetonitrile containing 0.1% formic acid (B), using the following elution profile: 0– 50 min, linear gradient 5–95% solvent B; 50–54 min, constant 95% solvent B; 54–60 min, constant 5% solvent B. Mass analysis was carried out in ESI positive ion mode (ESI^+^) and mass spectrometry conditions were as follows: spray voltage was set at 3.5 kV; source gases were set (in arbitrary units/min) for sheath gas, auxiliary gas and sweep gas at 40, 8, and 1, respectively; vaporizer temperature and ion transfer tube temperature were set at 320 °C and 275 °C, respectively. Survey MS scans of precursors were performed from 150 to 2,000 *m*/*z*, at 60 K resolution (full width of the peak at its half maximum, at 200 *m*/*z*) with MS parameters as follows: RF-lens, 50%; maximum injection time, 50 ms; data type, profile; internal mass calibration EASY-IC ^TM^ activated; AGC target: custom; normalized AGC target: 50%. A top speed (0.6 sec) data dependent MS^2^ was performed by isolation at 1.6 Th with the quadrupole, HCD fragmentation at a fixed 30% collision energy and analysis in the orbitrap at 30 K resolution (high resolution MS/MS analysis). The dynamic exclusion duration was set to 2.5 s with a 10 ppm tolerance around the selected precursor (isotopes excluded). Other MS/MS parameters were as follows: data type, profile; standard AGC target, internal mass calibration EASY-IC^TM^ activated. Mass spectrometer calibration was performed using the Pierce FlexMix calibration solution (Thermo Scientific). MS data acquisition was carried out utilizing the Xcalibur v. 3.0 software (Thermo Scientific).

### Generation of the molecular network and interpretation of MS/MS data

The raw data obtained from the LC-MS/MS system were converted to mzXML format using the ProteoWizard tool msconvert. All mzXML data were uploaded to the Global Natural Products Social Molecular Networking-platform (GNPS) and analyzed using the workflow as previously published (27), generating a molecular network based on the fragmentation spectra. To create consensus spectra, data were clustered by applying a parent mass tolerance of 1.0 Da and an MS/MS fragment ion tolerance of 0.5 Da; the remaining parameters were set to default. The output of the molecular networks was visualized using Cytoscape (51) and displayed using the settings “preferred layout” with “directed” style. For compounds in the network with known structures, e.g., DFO-B and acyl-DFOs, identified with GNPS IDs, the fragments were assigned to the corresponding molecules by comparison to the published fragmentation patterns. Structure analysis of compounds not giving rise to GNPS hits but linked to known compounds, was based on detailed comparison of MS/MS fragment spectra in order to identify both common and unique fragments.

## Acknowledgements

We acknowledge financial support from the IMPACT Biomolecules project of the Lorraine Université d’Excellence (Investissements d’avenir -ANR 15-004 to L.S., Y.S., K.J.W. and B.A.). We thank Prof. Helge B Bode from the Max Planck Institute for Terrestrial Microbiology, Marburg, Germany for his advice and support and our colleague Dr Christophe Jacob (UMR UL-CNRS IMoPA) for the valuable discussions on the project. We are also grateful to the Structural and Metabolomics Analyses Platform for the LC-MS/MS analyses (PASM, SF4242, Université de Lorraine, France) for access to the Orbitrap ID-X Tribrid Mass Spectrometer.

